# Bordetella Colonization Factor A (BcfA) elicits protective immunity against *Bordetella bronchiseptica* in the absence of an additional adjuvant

**DOI:** 10.1101/692830

**Authors:** Kacy S. Yount, Jamie Jennings-Gee, Kyle Caution, Audra R. Fullen, Kara N. Corps, Sally Quataert, Rajendar Deora, Purnima Dubey

**Affiliations:** Department of Microbial Infection & Immunity, The Ohio State University; Department of Microbiology & Immunology, Wake Forest University Health Sciences; College of Veterinary Medicine, Department of Veterinary Biosciences, The Ohio State University; Human Immunology Center Laboratory, University of Rochester Medical Center; Department of Microbiology, The Ohio State University

## Abstract

*Bordetella bronchiseptica (B. bronchiseptica)* is an etiologic agent of respiratory diseases in animals and humans. Despite widespread use of veterinary *B. bronchiseptica* vaccines, there is limited information on their composition, relative efficacy, and the immune responses they elicit. Furthermore, human *B. bronchiseptica* vaccines are not available. We leveraged the dual antigenic and adjuvant functions of BcfA to develop acellular *B. bronchiseptica* vaccines in the absence of an additional adjuvant. Balb/c mice immunized with BcfA alone or a trivalent vaccine containing BcfA and the *Bordetella* antigens FHA and Prn were equally protected against challenge with a prototype *B. bronchiseptica* strain. The trivalent vaccine protected mice significantly better than the canine vaccine Bronchicine^®^ and provided protection against a *B. bronchiseptica* strain isolated from a dog with kennel cough. Th1/17-polarized immune responses correlate with long-lasting protection against *Bordetellae* and other respiratory pathogens. Notably, BcfA strongly attenuated the Th2 responses elicited by FHA/Prn, resulting in Th1/17-skewed responses in inherently Th2-skewed Balb/c mice. Thus, BcfA functions as both an antigen and an adjuvant, providing protection as a single component vaccine. BcfA-adjuvanted vaccines may improve the efficacy and durability of vaccines against *Bordetellae* and other pathogens.

## INTRODUCTION

*Bordetella bronchiseptica (B. bronchiseptica)* is an animal pathogen with a wide host range, infecting farm and companion animals(1–6). It is one of the etiologic agents of kennel cough, or canine infectious respiratory disease (CIRD)(7). *B. bronchiseptica* is also increasingly isolated from immunocompromised humans such as those with HIV/AIDS, cancer, or cystic fibrosis. In many of these cases, the infections are linked to exposure to pets with *B. bronchiseptica*(8–10).

A nasal live attenuated(11) and a parenteral cellular antigen extract (CAe) vaccine (Bronchicine)(12) against *B. bronchiseptica* are widely used to minimize kennel cough outbreaks. The CAe formulation replaced more reactogenic whole cell inactivated vaccines in parallel to the development of acellular pertussis vaccines (aPV). However, a human vaccine against *B. bronchiseptica* is not available. Although antigens expressed by *B. bronchiseptica* are present in aPV against the human pathogen, *B. pertussis*(13, 14), these vaccines are only partially effective against *B. bronchiseptica*(15).

While considerable efforts have been devoted to evaluation of the immune response and effectiveness of aPV, there is insufficient research to determine the effectiveness of CAe *B. bronchiseptica* vaccines. Dogs vaccinated with CAe produced serum IgG and IgA and had reduced bacterial burden compared to unvaccinated dogs(16, 17). However, minor vaccine-related side effects were observed and coughing in 20% of immunized animals was reported, suggesting that the vaccine does not provide complete protection against disease(17). Furthermore, information on the immune response and protective efficacy of these vaccines is limited(12). Thus, there is an urgent need for well-defined, immunogenic acellular vaccines against *B. bronchiseptica* for veterinary and human use.

Together, Th1/17 cellular responses and Th1-skewed antibody responses provide long-lasting protective immunity against *Bordetellae*(18). At present, all aPV are adjuvanted with alum(13, 14), which elicits Th2-skewed cellular and humoral responses with sub-optimal and short-lived protection(18, 19). While alum does not cause pyrexia and has the strongest safety record of any adjuvant used in human vaccines(20), there have been reports of adverse reactions in animals and humans(21, 22). Thus, development of improved adjuvants is a pressing objective for more effective control of both veterinary and human diseases.

We previously reported identification of Bordetella colonization factor A (BcfA), an outer membrane protein expressed by *B. bronchiseptica* but not by the human pathogen, *B. pertussis*(23). BcfA is a paralog of outer membrane protein BipA and has significant homology to intimins and invasins of other bacteria(23). We showed that an experimental vaccine containing BcfA adsorbed to alum elicited protective immune responses against *B. bronchiseptica*(24).

BcfA is also an adjuvant that elicits Th1/Th17 cytokine responses and Th1-type antibodies to protein antigens(25) potentially serving as an alternative adjuvant to alum. In the present study, we tested the efficacy of BcfA as a monovalent vaccine and combined with *Bordetella* virulence factors FHA and Prn. We found that Th2-prone Balb/c mice immunized with BcfA as an antigen and without an additional adjuvant elicited Th1/17-polarized responses and efficiently cleared a *B. bronchiseptica* infection from the lungs and trachea. A combination vaccine containing BcfA and two *Bordetella* proteins FHA and Prn(14) also provided protection against laboratory and canine isolates of *B. bronchiseptica*. Protection by the BcfA-containing vaccine was superior to that provided by a current veterinary CAe vaccine. Together, our data show that the adjuvant and antigenic properties of BcfA will elicit highly protective immune responses against *B. bronchiseptica* for veterinary and human applications. Additionally, BcfA can function as an adjuvant to enhance immune responses against pathogens for which Th1/Th17 immune responses correlate with better protection(26, 27).

## RESULTS

### Immunization with BcfA as a single antigen in the absence of another adjuvant reduces *B. bronchiseptica* colonization of the mouse respiratory tract

We previously reported that immunization with BcfA/Alum protected mice against *B. bronchiseptica* challenge(24). BcfA also enhanced immune responses to heterologous antigens and to *Bordetella* vaccine antigens FHA and Prn(25). These results suggested a dual protective function of BcfA as an antigen and an adjuvant. Here, we first tested the hypothesis that BcfA as the sole component would protect against *B. bronchiseptica* infection in the absence of alum. Balb/c mice (male and female) were immunized intramuscularly (i.m.) with BcfA/Alum or BcfA alone (as described in Methods), and challenged with the prototype *B. bronchiseptica* laboratory strain RB50 (originally isolated from a rabbit)(28). CFUs in the lungs and trachea were enumerated at 4 days post-infection (dpi). Both immunizations protected the lungs and trachea of mice compared to naïve unimmunized mice. Bacterial burden was similar in both organs from mice immunized with BcfA/Alum or BcfA alone (Fig 1), demonstrating that alum is dispensable for protection mediated by BcfA.

**Figure 1.**
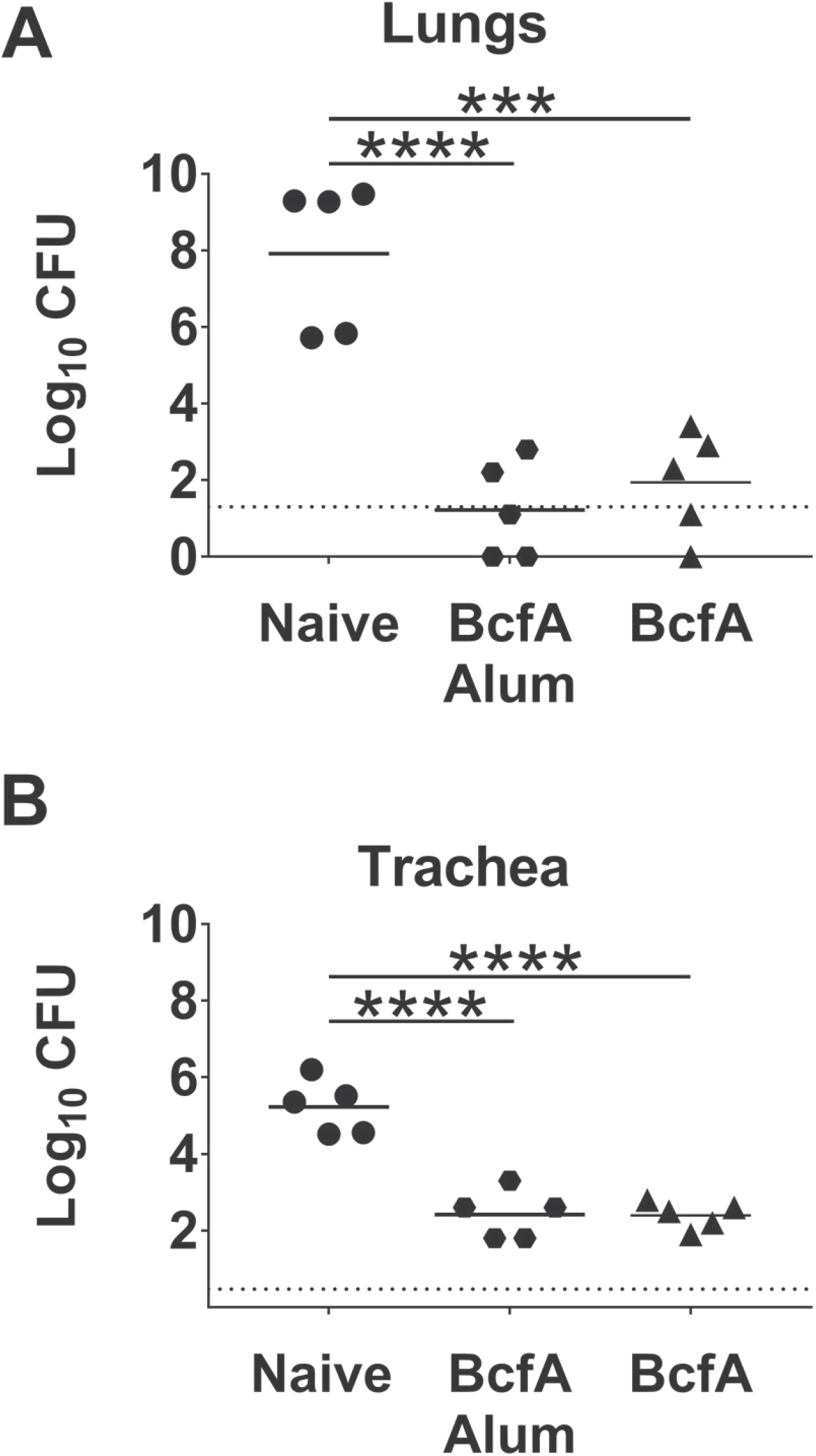
Immunization with BcfA alone is as effective as BcfA/Alum to reduce *B. bronchiseptica* colonization of the lungs and trachea. Balb/c mice (N=5/group) were immunized, challenged with RB50, and sacrificed at 4 dpi. Lungs and trachea were homogenized, serially diluted, and plated for CFU enumeration. (A) Lung CFUs. (B) Trachea CFUs. NS = not significant.

### A trivalent BcfA-adjuvanted vaccine is highly protective against *B. bronchiseptica*

FHA and Prn are *Bordetella* proteins that are antigens in aPV(13, 14) and have roles in adherence and pathogenesis of the human and animal pathogens(29–31). We tested whether a trivalent vaccine (BcfA/FHA/Prn) would provide superior protection compared to BcfA alone. Balb/c mice immunized with BcfA alone, FHA/Prn, or BcfA/FHA/Prn were challenged with RB50. CFUs were enumerated from lungs and trachea at 3 or 7 dpi. At both time points, the lungs (Fig 2A, 2B) and trachea (Fig 2C, 2D) of naïve challenged mice were highly colonized by RB50. Compared to FHA/Prn-immunized mice, BcfA-immunized mice had significantly reduced bacterial burden in the lungs at 3 dpi, with CFUs at or below the limit of detection (20 CFUs) in 4 out of 8 mice (Fig 2A). At 7 dpi, compared to naïve challenged mice, both FHA/Prn- and BcfA-immunized mice exhibited reduced bacterial burden in the lungs (FIG 2B) and trachea (Fig 2D).

**Figure 2.**
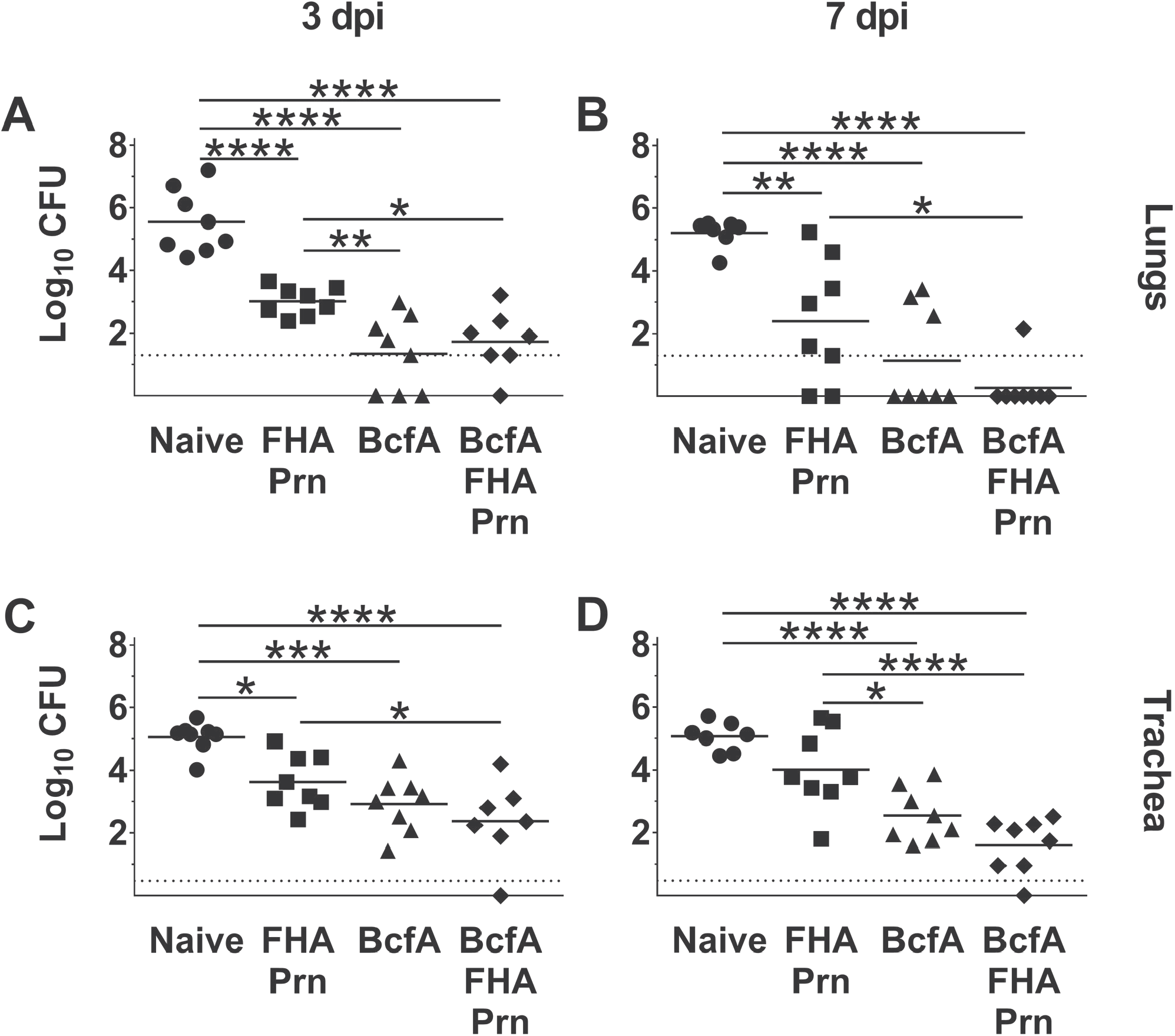
Immunization of mice with a monovalent BcfA vaccine or a trivalent vaccine with FHA and Prn reduces *B. bronchiseptica* bacterial burden from the lungs and trache. Balb/c mice (N=8/group) were unimmunized (naïve) or immunized with FHA/Prn, BcfA, or BcfA/FHA/Prn, and challenged with RB50. Lung CFUs at (A) 3 dpi and (B) 7 dpi. Trachea CFUs at (C) 3 dpi and (D) 7 dpi. One representative experiment of 2 is shown. * P < 0.05, ** P < 0.01, *** P < 0.001, **** P < 0.0001

While one mouse immunized with BcfA/FHA/Prn had no detectable bacteria in the lungs (Fig 2A) or trachea (Fig 2C) at 3 dpi, the average bacterial load was not significantly different than FHA/Prn- or BcfA-immunized mice. At 7 dpi, BcfA/FHA/Prn immunization was significantly better than FHA/Prn, with ~2 log_10_ lower bacterial load in both organs (Fig 2B, 2D). Strikingly, there was no statistical difference between the bacterial burdens of BcfA/FHA/Prn- and BcfA-immunized mice. Together, these data show that BcfA alone, without an additional adjuvant, and a trivalent BcfA-containing vaccine reduce bacterial load in the lungs and trachea.

### Immunization with BcfA alone elicits an antibody response of similar magnitude but with a more pronounced Th1-skewed phenotype compared to BcfA/Alum

We previously reported that BcfA/Alum immunization elicited BcfA-specific IgG2 antibodies in C57BL/6 mice(24). This observation was noteworthy because alum-adjuvanted vaccines including aPV elicit Th2-polarized responses(18, 32–34) and because Th1/17, but not Th2, responses are critical for immunity against *Bordetellae*(18). We evaluated BcfA-specific antibodies in the serum of mice immunized with BcfA/Alum, BcfA, or BcfA/FHA/Prn. All three immunizations produced a similar level of total IgG in the serum (Fig 3A) and lungs (Fig 3B).

**Figure 3.**
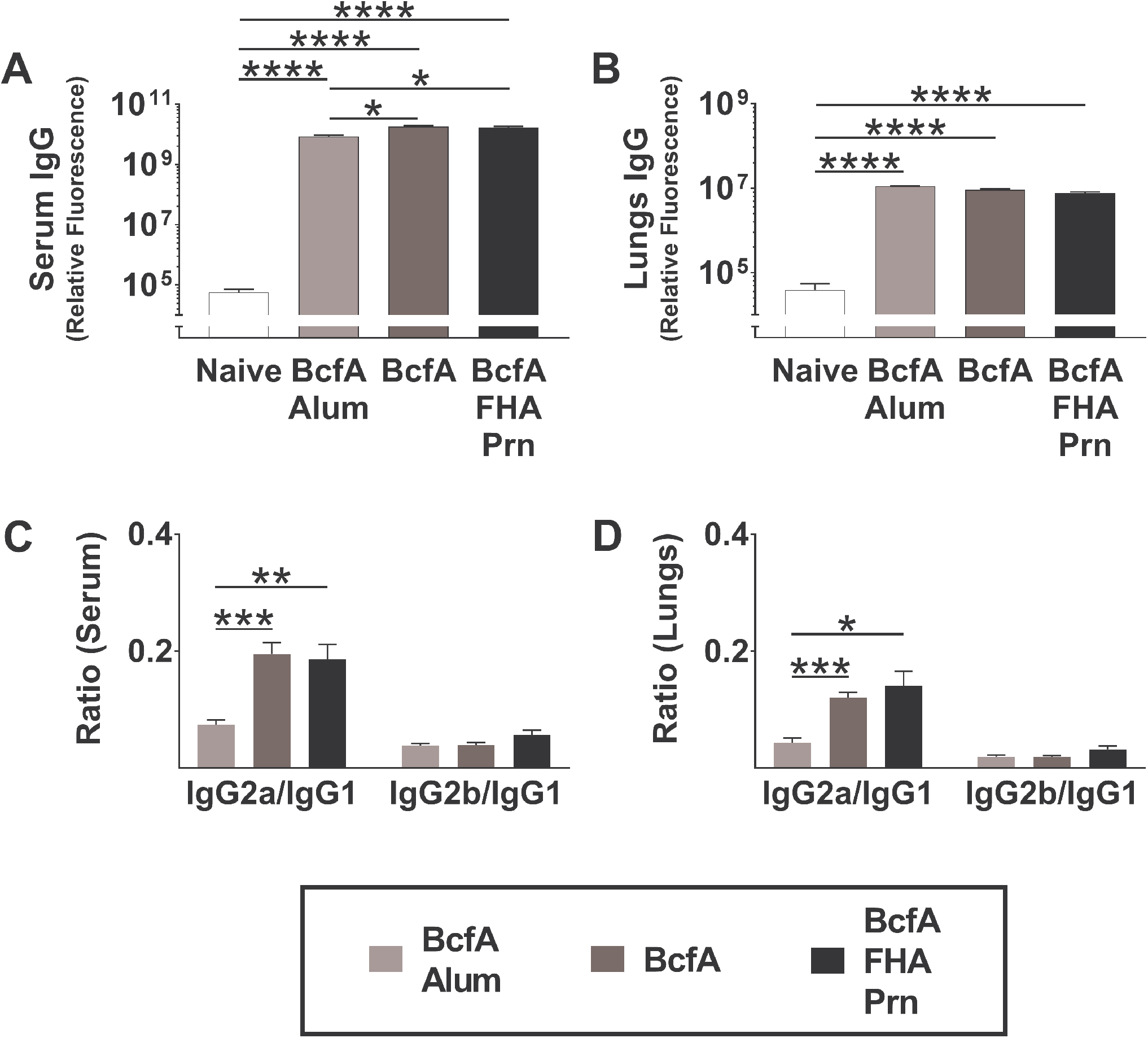
Immunization with BcfA alone or with BcfA/FHA/Prn elicits strong systemic and mucosal BcfA-specific Th1-type antibody responses. BcfA-specific total IgG as well as antigen-specific isotypes IgG1, IgG2a, and IgG2b were quantified in serum and lung homogenates at 3-4 dpi by multiplex fluorescent assay (N=5-8/group). Results are log_10_-transformed and presented as relative fluorescence units with background subtracted. (A) BcfA-specific IgG in serum. (B) BcfA-specific IgG in lung homogenate. (C) BcfA-specific IgG2/IgG1 isotype ratios in serum. (D) BcfA-specific IgG2/IgG1 isotype ratios in lungs. * P < 0.05, ** P < 0.01, *** P < 0.001, **** P < 0.0001

We observed a higher ratio of BcfA-specific IgG2a/IgG1 in the serum (Fig 3C) and lungs (Fig 3D) of mice immunized with BcfA alone or with BcfA/FHA/Prn compared to with BcfA/Alum. Thus, removing alum from the vaccine reduces the Th2-type of BcfA-specific antibodies while maintaining the magnitude of the IgG response. In addition, a higher ratio of IgG2/IgG1 antibodies suggests a Th1-polarized response that is correlated with better protection against *Bordetella* infection(18, 35, 36).

### BcfA elicits Th1-type antibody responses to FHA

FHA and Prn alone or adjuvanted to alum elicit Th2-skewed antibody responses(25). To determine whether BcfA remodeled these responses towards Th1 we evaluated the systemic and mucosal IgG levels and calculated the ratio of FHA- and Prn-specific IgG2/IgG1 antibodies elicited by BcfA/FHA/Prn. FHA-specific IgG was higher in the serum (Fig 4A) and lungs (Fig 4C) of FHA/Prn- and BcfA/FHA/Prn-immunized mice compared to naïve mice, while Prn-specific antibody levels were increased in the serum (Fig 4B), but not the lungs (Fig 4D). We observed a higher ratio of FHA-specific IgG2b/IgG1 in the serum (Fig 4E) and lungs (Fig 4G) of mice immunized with BcfA/FHA/Prn compared to FHA/Prn alone. These data suggest that the adjuvant function of BcfA shifts the antibody response to FHA toward Th1 by reducing the Th2-type antibodies. In contrast, the Prn-specific antibody ratios were not altered (FIG 4F).

**Figure 4.**
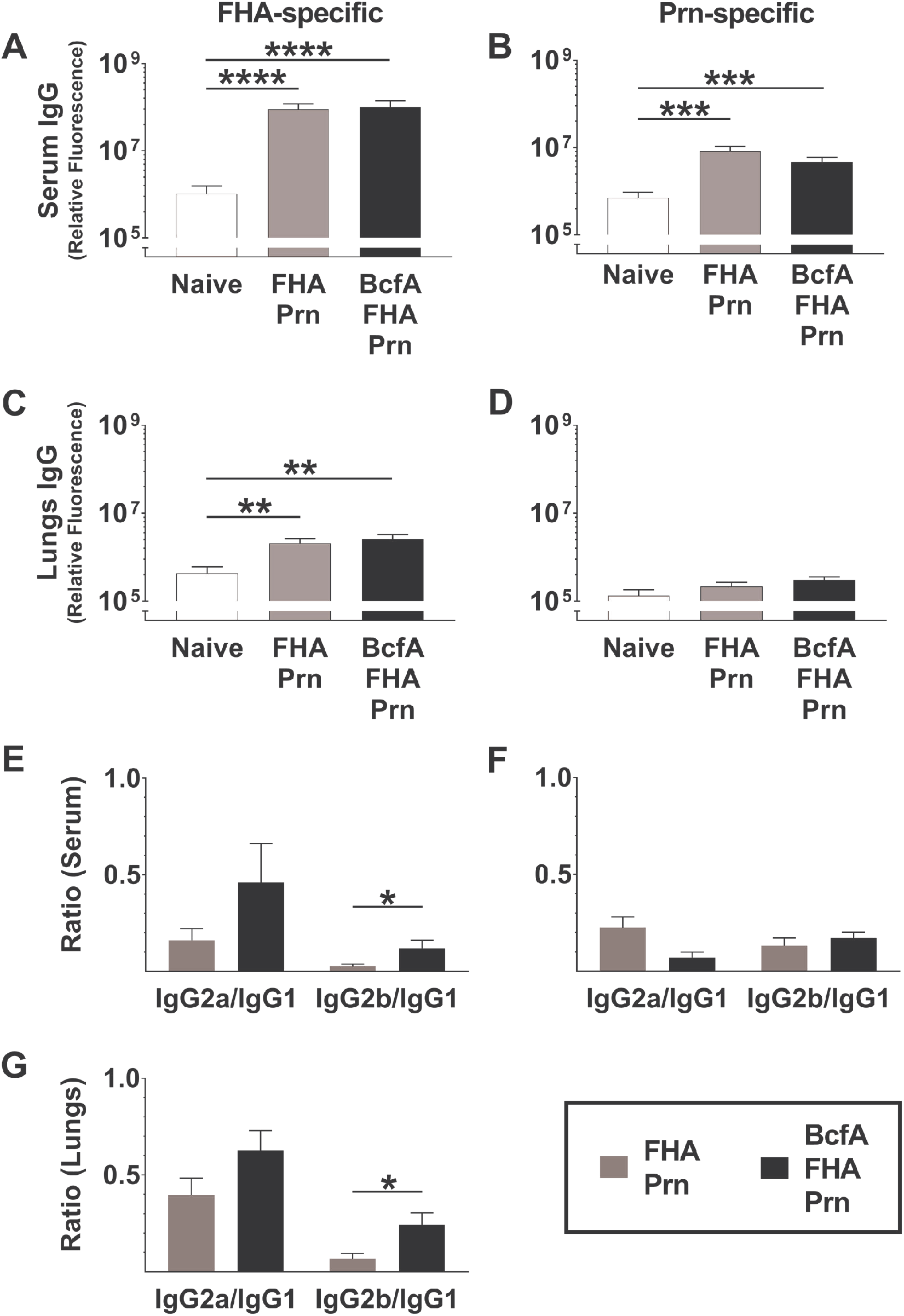
The addition of BcfA to FHA/Prn immunization elicits Th1-type FHA- and Prn-specific antibody responses. FHA- and Prn-specific total IgG and isotypes IgG1, IgG2a, and IgG2b were quantified in serum and lung homogenates at 3-4 dpi by multiplex fluorescent assay (N=8/group). Results are log_10_-transformed and presented as relative fluorescence units with background subtracted. (A) FHA-specific IgG in serum. (B) Prn-specific IgG in serum. (C) FHA-specific IgG in lung homogenate. (D) Prn-specific IgG in lung homogenate. (E) FHA-specific IgG2/IgG1 isotype ratios in serum. (F) Prn-specific IgG2/IgG1 isotype ratios in serum. (G) FHA-specific IgG2/IgG1 isotype ratios in lung homogenate. * P < 0.05, ** P < 0.01

### Immunization with BcfA/FHA/Prn elicits Th1/17 cytokine production and attenuates Th2 cytokine production

We showed that murine bone marrow dendritic cells stimulated with BcfA produced Th1/17 polarizing innate cytokines including IL-12/23. Furthermore, the addition of BcfA to aPV elicited Th1/17-polarized immune responses in C57BL/6 mice by attenuating the Th2 cytokine responses observed with alum-adjuvanted aPV(25). C57BL/6 mice have an inherently Th1-skewed immune phenotype(37). Here, we tested whether immunization of the Th2-prone Balb/c mouse strain(37) with a BcfA-adjuvanted vaccine would similarly shift T cell responses towards Th1/17.

To evaluate systemic responses, splenocytes from FHA/Prn- and BcfA/FHA/Prn-immunized mice were stimulated *in vitro* with FHA, Prn, or BcfA for 7 days, and quantified cytokines present in the supernatants by ELISA. Splenocytes from BcfA/FHA/Prn immunized mice produced significantly lower amounts of Th2 cytokines IL-5 (Fig 5A) and IL-13 (Fig 5B) compared to FHA/Prn-immunized spleen cells. Similar levels of FHA- and Prn-specific Th1 effector cytokine IFNγ (Fig 5C) and Th17 effector cytokine IL-17 (Fig 5D) were produced by both immunizations while high levels of BcfA-specific IFNγ and IL-17 (FIG 5C, 5D) were produced from spleens of BcfA/FHA/Prn-immunized mice. Together, these results show that, by inhibiting the production of Th2 cytokines, BcfA skews responses away from Th2 and toward Th1/Th17 in a Th2-prone mouse strain.

**Figure 5.**
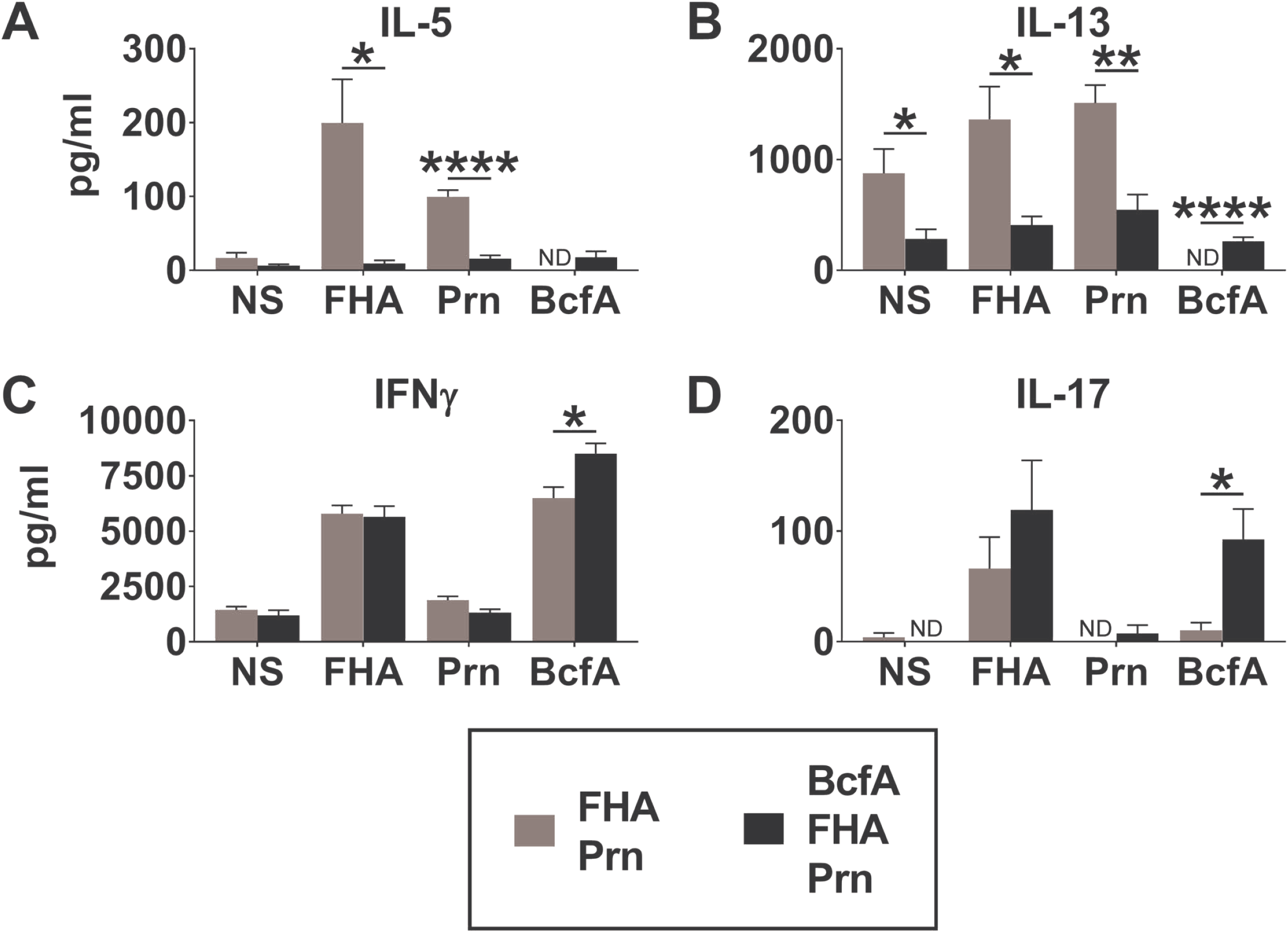
BcfA/FHA/Prn immunization attenuates Th2 cytokines IL-5 and IL-13 compared to FHA/Prn immunization. Splenocytes harvested from Balb/C mice (N=8/group) at 3 dpi were stimulated *in vitro* with medium alone (NS) or with 1 μg/ml FHA, Prn, or BcfA for 7 days. (A) IL-5, (B) IL-13, (C) IFNγ, and (D) IL-17 in culture supernatants were quantified by ELISA. * P < 0.05, ** P < 0.01, **** P < 0.0001

### BcfA/FHA/Prn provides better protection against a laboratory and a clinical strain of *B. bronchiseptica* than a commercial cellular antigen extract vaccine

We compared protection provided by BcfA or BcfA/FHA/Prn to that provided by Bronchicine^®^, a widely used but insufficiently characterized veterinary vaccine. Mice were immunized with BcfA/FHA/Prn or BcfA as above or with 1/10^th^ or 1/5^th^ canine dose of Bronchicine^®^, doses similar to those of human aPV commonly tested in mice(25, 38). Immunized and naïve mice were subsequently challenged with RB50 or with MBORD 685, a canine *B. bronchiseptica* strain isolated from a dog with kennel cough(3). Overall, colonization of the respiratory tract of naïve and immunized mice by MBORD 685 was equivalent to colonization by the rabbit isolate RB50 (Fig 6A,B). While all four immunizations reduced bacterial burden compared to naïve mice, BcfA/FHA/Prn immunization most efficiently reduced bacterial burden in the lungs (Fig 6A) and trachea (Fig 6B). Immunization with BcfA alone was significantly more protective than 1/10^th^ but not 1/5^th^ dose Bronchicine^®^ in both the lungs (Fig 6A) and trachea (Fig 6B). Importantly, both doses of Bronchicine^®^ were significantly less protective in the lungs of immunized mice than BcfA/FHA/Prn (Fig 6A). The lungs of 83% of mice immunized with BcfA/FHA/Prn were cleared of *B. bronchiseptica* below the limit of detection while only - 23% and 50% were cleared by 1/10 and 1/5 dose Bronchicine^®^, respectively (Fig 6A). Together, these results support the clinical applicability of either monovalent BcfA or trivalent BcfA/FHA/Prn as veterinary vaccines and provide a new avenue for more effective and, potentially, durable protection against this pathogen.

**Figure 6.**
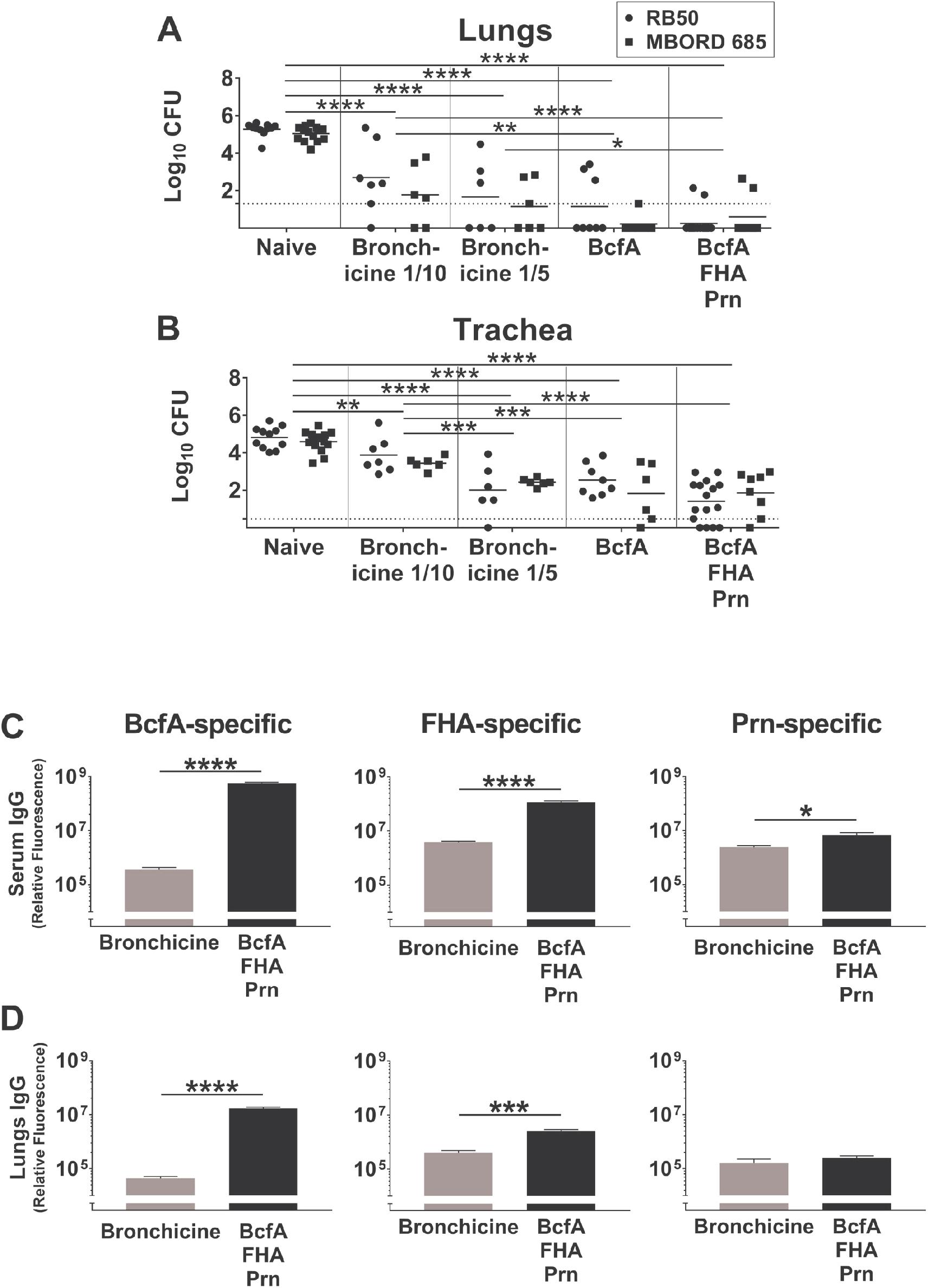
BcfA/FHA/Prn immunization is more effective and elicits a stronger antibody response than Bronchicine^®^. Balb/c mice (N=6-16/group) were immunized and challenged with RB50 or MBORD 685. CFUs from (A) lungs and (B) trachea on 7dpi. Antigen-specific total IgG in (C) serum and (D) lung homogenate. Data are log_10_-transformed relative fluorescence units with background subtracted. * P < 0.05, ** P < 0.01, *** P < 0.001, **** P < 0.0001

### BcfA/FHA/Prn elicits more robust antigen-specific antibody responses than a commercial veterinary vaccine

We observed similar FHA- and Prn-specific antibody levels between 1/10 Bronchicine^®^-immunized mice and naïve mice (see Fig 4). In contrast, immunization with BcfA/FHA/Prn elicited higher serum antibody responses to all three antigens (Fig 6C) and lung antibody responses to BcfA and FHA (Fig 6D). Together, these results show that BcfA/FHA/Prn elicits a stronger immune response and is more protective than Bronchicine^®^ in a murine model of *B. bronchiseptica* infection.

### Immunization reduces inflammation in the lungs of mice challenged with *B. bronchiseptica*

*B. bronchiseptica* infection of unimmunized mice causes considerable damage to lung tissues(24, 39). We determined whether immunization decreased lung injury compared to unimmunized mice. Blinded H&E sections were evaluated for several parameters of lung injury and immune cell infiltration (Supplemental Table 1). As expected, total pathology score for naïve challenged mice (Fig 7A) was highest, exhibiting severe degeneration and airway necrosis that resulted in airway obliteration, markedly thickened alveolar walls and considerable influx of viable and degenerate polymorphonuclear cells (PMNs).

**Figure 7.**
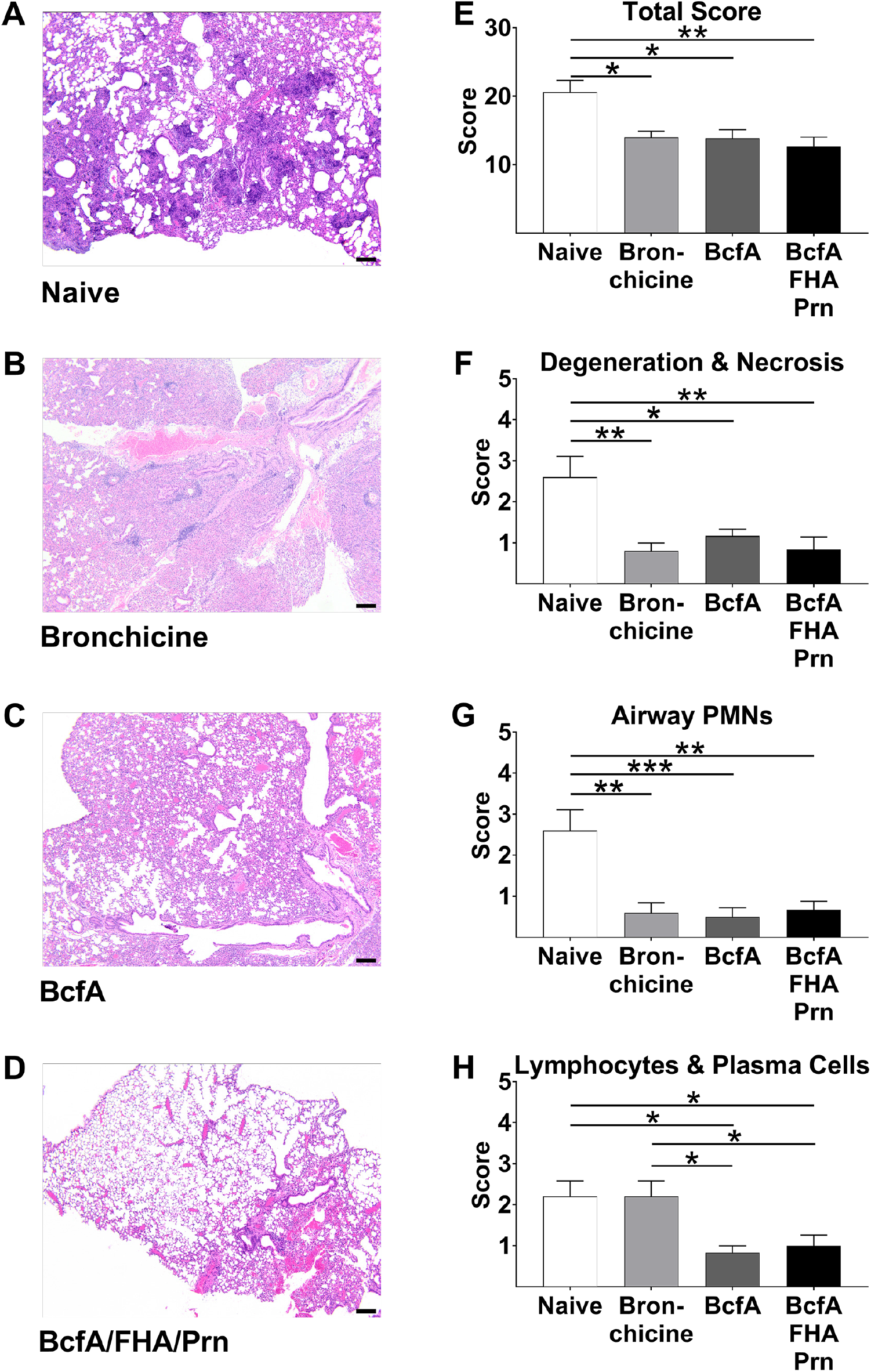
BcfA-containing vaccines and Bronchicine^®^ reduce lung injury and elicit characteristically different cell infiltrates. Balb/c mice (N=5-6/group) were immunized, challenged with RB50, and sacrificed 3-4 dpi. H&E stained lung sections (A-D) were scored qualitatively from 0-5 for (E) total pathology score, (F) degeneration and necrosis, (G) infiltrating airway PMNs, and (H) infiltrating lymphocytes and plasma cells. Scale bar in A-D, 200 μm. * P < 0.05, ** P < 0.01, *** P < 0.001

In contrast, immunized mice (Fig 7B-D) had significantly lower pathology scores compared to naïve mice (Fig 7E), as well as reduced degeneration/necrosis (Fig 7F) and infiltrating airway PMNs (Fig 7G). Bronchicine^®^(1/10 dose)-immunized mice exhibited significantly more infiltrating lymphocytes and plasma cells (Fig 7H) compared to lungs of BcfA and BcfA/FHA/Prn-immunized animals. Thus, there are qualitative differences between highly effective BcfA-containing vaccines and the poorly protective Bronchicine^®^.

## DISCUSSION

There is an urgent need for improved vaccines against *B. bronchiseptica* since respiratory diseases caused by this pathogen are a significant health concern for animals and humans. Although vaccines are widely used in veterinary medicine to prevent kennel cough in dogs, information regarding their composition, immune profile or protective efficacy is sparse. An effective vaccine must contain protective antigens that elicit strong immune responses and adjuvants that heighten the response and elicit immune phenotypes that reflect the responses generated by natural infection. Cellular Th1/17 responses and humoral Th1-skewed responses generated by whole cell vaccines or natural infection are required for long-lived protection against *Bordetellae* (18) and other pathogens (26, 27).

Though alum has been used successfully as a vaccine adjuvant since the early 1900s to prevent disease, it elicits Th2-skewed responses and thus weaker and shorter-lived immunity(40). Thus, substituting alum with Th1/17 polarizing adjuvants is likely to improve vaccine-induced immunity. We previously showed that BcfA elicits Th1/17 responses in C57BL/6 mice(25). To determine the full potential of BcfA in the absence of alum to mediate protective shifts in immune responses, we administered BcfA-containing vaccines to the Th2-prone mouse strain, Balb/c(37). Cellular and humoral responses to BcfA, FHA, and Prn were remodeled to Th1/17, primarily by attenuating Th2 cytokine production. This also resulted in a higher ratio of IgG2/IgG1 antibodies. Together, these data provide further evidence of the adjuvant activity and immune modulatory functions of BcfA.

Animals immunized with BcfA alone reduced RB50 bacterial CFUs as effectively as mice immunized with BcfA/Alum, demonstrating that BcfA is a protective antigen and does not require an additional adjuvant. It is striking that a single protein is a strong antigen and adjuvant. Conversely, other well-characterized *Bordetella* virulence factors such as FHA(41, 42), adenylate cyclase toxin (ACT)(43), lipooligosaccharide (LOS)(44), and BopN(45) shift the cellular immune response away from Th1 and/or toward Th2. Thus, it is notable that the BcfA-containing vaccine characterized in this study attenuates the Th2 responses elicited by FHA.

We hypothesized that the trivalent BcfA/FHA/Prn vaccine would be more effective than BcfA alone due to the presence of two additional antigens. Surprisingly, the protection provided by this combination was not significantly better than BcfA alone, although cytokine and antibody responses to FHA were detected. We did not detect strong responses to Prn, suggesting that this antigen may be dispensable. Furthermore, allelic variants of Prn in *B. bronchiseptica* are reported(46, 47), implying that Prn-specific responses may not be protective due to antigenic drift among circulating strains. We showed previously that *B. bronchiseptica* isolates from dogs (including MBORD 685 used in this study), cats, horses, pigs, and humans highly express BcfA (10, 24). Production of BcfA by strains isolated from companion and food-producing animals strengthens its utility as a protective antigen in a novel vaccine, a possibility supported by our data showing that both BcfA alone and BcfA/FHA/Prn immunizations reduced MBORD 685 bacterial CFUs.

To determine the relative efficacy of our acellular formulations to currently used veterinary vaccines, we compared the protection provided by BcfA/FHA/Prn to Bronchicine^®^. FHA and Prn have been detected at low levels in Bronchicine^®^(16), and we detected low levels of BcfA (data not shown). Thus, despite shared antigenicity, BcfA/FHA/Prn immunization elicited stronger responses and provided superior protection compared to this current veterinary vaccine. Differences in antigen quantity (unknown in Bronchicine^®^) may, at least in part, explain the difference in protection. In addition, Bronchicine^®^ elicits weaker antibody responses than previous whole cell bacterin vaccines, likely due to the reduction of LOS, which adjuvants immune responses but is also reactogenic(12). Furthermore, CAe formulations may present inhibitory proteins or polysaccharides that attenuate effector responses(48, 49). Differences in the composition and volume of immune cell infiltration elicited by the CAe or BcfA-containing vaccines may also contribute to varied protection.

Together, our data suggest that an acellular component vaccine, leveraging the dual antigenic and adjuvant function of BcfA as a monovalent or trivalent vaccine formulation, has strong potential as a novel immunization approach for animal and human respiratory diseases mediated by *B. bronchiseptica*. Furthermore, the adjuvant function of BcfA may improve immunity against other bacterial and viral pathogens that require Th1/17 responses for protection against disease(26, 27).

## MATERIALS AND METHODS

### Bacterial strains, media, and growth conditions

Wild-type *B. bronchiseptica* strain RB50(28), and canine isolate MBORD 685(3), were maintained on Bordet-Gengou (BG) agar (Difco) containing 7.5% defibrinated sheep’s blood supplemented with 100 μg/ml streptomycin. For animal inoculations, liquid cultures from single colonies were grown at 37°C on a roller drum to OD_600_ ≈ 1.0 in Stainer-Scholte medium and 100 μg/ml streptomycin.

### Animals

All experiments were reviewed and approved by the Ohio State University Institutional Animal Care and Use Committee (Protocol #2017A00000090). Balb/c mice (male and female, 6 to 21 weeks old) were bred in-house.

### Reagents

FHA and Prn derived from *B. pertussis* were purchased from Kaketsuken (Japan) and List Biologicals (Campbell, CA), respectively. BcfA was produced and purified as described previously(23). Endotoxin levels in all proteins were at acceptable levels and below that of aPV(50). Bronchicine^®^ CAe (Zoetis) was purchased from OSU Veterinary Biosciences pharmacy. RPMI was from Thermo Fisher Scientific (Waltham, MA). Fetal bovine serum (FBS) was from Sigma-Aldrich (St. Louis, MO). ELISA kits were from eBioscience (Thermo Fisher Scientific).

### Immunizations

Mice were lightly anesthetized with 2.5% isoflurane–O_2_ for i.m immunization on day 0 and boost on day 28-35 as demonstrated previously(51) with the following vaccines: a) 1/10^th^ or 1/5^th^ canine dose of Bronchicine^^®^^ or b) 100 μl of experimental acellular vaccine containing varying combinations of 1.6 μg FHA, 0.5 μg Prn, and 30 μg BcfA. In alum-containing immunizations, 130 μg of aluminum hydroxide colloidal suspension (Sigma) was used.

### Bacterial challenge

Bacterial strains RB50 and MBORD685 grown overnight to OD_600_ ≈ 1.0 were diluted in PBS to 1×10^5^-5×10^5^ bacteria per 50 μl. On days 14-20 post-boost, mice were lightly anesthetized with 2.5% isoflurane–O_2_ and the 50 μl inoculum was delivered to both nares as demonstrated previously(51).

### Colony enumeration

Mice were euthanized at 3-7 dpi and the lungs, trachea, nasal septum, spleen, and blood were harvested as demonstrated previously(51). Respiratory tract tissues were mechanically disrupted in PBS + 1% casein and various dilutions were plated on BG agar containing 7.5% sheep’s blood and supplemented with 100 μg/ml streptomycin. Colony forming units (CFUs) were counted after 2 days of incubation at 37° C. Data were transformed to log_10_. Dotted line in each Figure indicates limit of detection at 20 CFUs for lungs and 3 CFUs for trachea.

### Splenocyte stimulation and ELISAs

Spleens were dissociated and red blood cells were lysed. Single cell suspension was plated at 2.5 × 10^6^ cells/well of complete T cell media (RPMI, 10% FBS, 10 μg/ml gentamicin, 5 × 10^-5^ M 2-mercaptoethanol) and stimulated with 1 μg/ml FHA, Prn, or BcfA or media alone as negative control. Supernatant was collected on day 7 post-stimulation. Production of IFNγ, IL-5, IL-13 and IL-17 was quantified by sandwich ELISA according to the manufacturer’s instructions.

### Antibody analysis

Purified antigens were coupled through an amine linkage to MagPlex C magnetic microspheres (Luminex Corporation), each with a unique fluorescent bead region address, and combined to form a 5-plex microarray. Mouse serum or lung homogenates were diluted in assay buffer, PBS–0.1% Brij-35–1% bovine serum albumin (BSA), pH 7.2, and incubated with the beads for 2 h at room temperature (r.t.) in the dark while shaking at 800 rpm. After washing, appropriate biotinylated detection antibody was added, i.e., goat anti-mouse total IgG, rat anti-IgG1, rat anti-IgG2a, rat anti-IgG2b, or goat anti-IgG2c, at a 1:250 dilution in assay buffer for 1 h at r.t. After washing, streptavidin-phycoerythrin (SA-PE) at 1:250 in assay buffer was added for 1 h with shaking. Unbound SA-PE was removed by washing, and the beads were resuspended in 100 μl PBS prior to reading on a Luminex 200 flow cytometer. Antibody isotype and subclass values are reported in arbitrary fluorescent intensity units. Antibody ratios were calculated by dividing the fluorescent intensity units of Ig2a or IgG2b by IgG1 after subtracting background and accounting for sample dilution.

### Histology and Scoring

The superior lobe of the right lung was harvested from mice 3-4 dpi and fixed in 2 ml of 10% neutral buffered formalin for at least 24 h. Tissues were processed, and embedded in paraffin. Five micron sections (3 per tissue) were stained with hematoxylin and eosin by the Comparative Pathology & Mouse Phenotyping Shared Resource at The Ohio State University. A board-certified veterinary pathologist (KNC) was blinded to experimental groups and sections were scored qualitatively 0-5 for degree of cellularity and consolidation, thickness of alveolar walls, degeneration and necrosis, edema, hemorrhage, infiltrating alveolar/interstitial polymorphonuclear cells (PMNs), intrabronchial PMNs, perivascular and peribronchial lymphocytes and plasma cells, and alveolar macrophages. Total inflammation score was calculated by totaling the qualitative assessments in each category.

### Statistical analysis

Bacterial CFUs and antibody levels were evaluated using a one-way analysis of variance (ANOVA) with Holm-Sidak correction for multiple comparisons for experiments with 3 or more groups and using student’s t-test for experiments with 2 groups. For grouped analyses of CFUs, two-way ANOVA with Holm-Sidak correction for multiple comparisons was used to compare immunization groups. Antibody ratios and cytokine levels were evaluated by multiple student’s t-tests with Holm-Sidak correction for multiple comparisons. Pathology scores were evaluated using a one-way ANOVA with Holm-Sidak correction for multiple comparisons.

## Acknowledgements

This work was supported by RO1AI125560, funds from Wake Forest Innovations, and start-up funds from The Ohio State University. We acknowledge Jennifer Bruno, Jessica Brown, Jessica Ferman, and Bhavneep Kaur for technical assistance, and the Comparative Pathology Core laboratory at OSU for tissue processing. We thank Eugene Oltz for critical review of the manuscript.

## Author Contributions

KSY, PD, and RD designed experiments. KSY, JJ-G, KC and AF conducted experiments. SQ conducted antibody analysis and KNC conducted histology analysis. KSY, PD, and RD interpreted data and wrote the manuscript.

